# Linking time-lagged functional dynamics to spatial constraints in resting-state fMRI

**DOI:** 10.64898/2026.05.24.727506

**Authors:** Danilo Benozzo

**Affiliations:** Department of Brain and Behavioral Sciences, University of Pavia, Pavia, Italy

**Keywords:** resting-state fMRI, brain dynamics, linear state-space model, oscillatory modes, distance matrix

## Abstract

Linear state-space models have been shown to effectively reproduce large-scale brain dynamics. We applied this approach to resting-state fMRI data acquired from 20 mice, focusing on the system’s Jacobian matrix, i.e. the effective connectivity, and specifically on its component encoding nonzero-lag interactions: the differential covariance matrix. Within this matrix, we concentrated on the off-diagonal component (dC-Cov), which reflect endogenous time-lagged correlations. Our aim was to identify a decomposition of the Jacobian matrix that facilitates its interpretation from a mechanistic perspective. Since the dC-Cov captures the rotational component of signal trajectories, we employed Schur decomposition to extract 2D rotational modes, each characterized by a pair of orthogonal vectors, and an associated angular frequency. This provides a more generative formulation of the modeling framework, thereby reducing the interpretability gap between this approach and connectome-based network models of coupled neural masses. Within this framework, the precision matrix governs the coupling between different Schur modes, while we hypothesize that the dC-Cov reflects spatial constraints imposed by inter-regional distances. By examining the relationship between dC-Cov and structural constraints imposed by the spatial placement of brain areas, we found a consistent alignment between the faster Schur modes across mice and the leading eigenvectors of the structural distance matrix.

## 1 Introduction

Resting-state functional magnetic resonance imaging (rs-fMRI) has become a central tool for studying large-scale brain dynamics. A common modeling strategy in this context relies on linear state-space models, which aim to infer latent dynamical interactions from observed brain activity [40, 45, 31]. This class of models is often referred to as a top-down or inverse problem approach, as the governing equations of the system are inferred directly from the data rather than specified a priori [14, 57]. Within this framework, the state-interaction matrix, i.e. the Jacobian of the dynamical system, is commonly interpreted as effective connectivity. Although the notion of effective connectivity originates from Dynamic Causal Modelling (DCM) [21], initially developed for task-based experiments, it has increasingly been adopted in the analysis of resting-state data [54, 23, 50].

While effective connectivity has a consolidated interpretation in task-based paradigms, its application to resting-state fMRI poses substantial challenges. Resting-state analyses typically involve an order of magnitude more brain regions than task-based studies, rendering hypothesis-driven strategies impractical. Moreover, element-wise interpretations of the Jacobian (where positive entries are labeled as excitatory and negative entries as inhibitory) become increasingly ambiguous as network size grows [54, 6]. As a result, both interpretability and scalability emerge as critical limitations of current top-down approaches when applied to fine-grained resting-state parcellations.

From a dynamical systems perspective, and under the assumption of linearity, the state-interaction matrix fully determines how brain activity evolves and propagates over time. A principled way to interpret this matrix is offered by its decomposition into the differential covariance and the precision matrix [35, 37, 9]. This decomposition separates time-lagged interactions from instantaneous statistical dependencies, providing a structured representation of the system dynamics that goes beyond element-wise inspection of the Jacobian.

In this work, we focus on the differential covariance due to its role in shaping the time-lagged pattern of interactions between brain regions [61] and in characterizing non-equilibrium dynamics [38]. In particular, its anti-symmetric component directly quantifies the violation of detailed balance and is proportional to the entropy production rate of the system [25]. A non-symmetric differential covariance therefore signals the presence of irreversible, i.e. out-of-equilibrium dynamics, an increasingly reported feature of macroscale brain activity [48, 34, 5].

A substantial body of work has characterized the temporal and spatial structure of fMRI signals through functional connectivity, primarily focusing on zero-lag correlations. By contrast, time-lagged synchronization has been less explored, largely because BOLD signals require careful disentanglement of phase delays originating from hemodynamic responses versus neural dynamics. Despite these challenges, early work demonstrated the presence of structured lagged correlations [41], with subsequent studies linking their spatial organization to arousal and vigilance states [53, 26]. These propagating patterns of lagged activity have been shown to align with the cortical hierarchical axis [68]. And from a modeling standpoint, their emergence has been parsimoniously explained by a combination of standing and traveling waves underlying spontaneous low-frequency brain fluctuations [8].

Here, we leveraged the Schur decomposition, an algebraic tool that allows the anti-symmetric differential covariance to be expressed as a combination of independent two-dimensional rotational modes. Each Schur mode is characterized by a specific angular frequency and a pair of orthogonal basis vectors, defining the coordinate plane on which the rotation occurs. This representation provides a natural interpretation of time-lagged interactions as oscillatory rotations unfolding in a low-dimensional subspace of the full state space. Building on this formulation, we investigated whether these rotational Schur modes are constrained by geometrical properties of the brain. Specifically, we examined their relationship with the spatial distance between brain regions defined by the parcellation used to cluster the empirical recordings.

Unlike structural connectivity, spatial distance is often overlooked in resting-state modeling, despite its relevance for signal propagation and conduction delays [39]. Structural strength and distance are tightly coupled through the exponential distance rule (EDR), whereby connectivity decays exponentially with inter-regional distance. Originally identified in macaque white matter [19], the EDR has since been confirmed across species and tissue types, including mouse gray and white matter [29, 30] and more recently in Drosophila, marmoset, and humans [33, 52]. This cross-species consistency suggests that distance encodes a fundamental geometric constraint balancing wiring cost and communication efficiency.

In parallel, several studies have investigated the relationship between spatial distance and functional organization in fMRI data. Early work reported a decay of functional correlations with distance, following an inverse-square law [58], along with a frequency-dependent modulation whereby low frequencies preferentially support long-distance intrahemispheric and bilaterally homologous regions [59]. More recently, a power-law relationship between functional correlation and distance has been interpreted as a signature of turbulent-like-dynamics [15]. The relevance of geometry in shaping brain dynamics has also been emerged from graph spectral approaches, where eigenmodes derived from distance-based adjacency matrices capture major features of fMRI activity, albeit less accurately than models incorporating cortical surface geometry [47]. Efforts to jointly integrate structure, distance, and function have further shown that embedding distance-derived delays into structural Laplacians improves the alignment between modeled dynamics and canonical resting-state networks [66].

Our analyses of a resting-state mouse dataset reveal that Schur modes associated with higher oscillatory frequencies, i.e. principal Schur modes, exhibit a consistent alignment with the dominant eigenvectors of the Laplacian matrix constructed from the distance matrix. And by approximating the asymmetric part of the differential covariance with a low-rank reconstruction based on the dominant eigenvectors of the distance-based Laplacian matrix we replicated better than chance both the signal power spectral density and imaginary part of coherence. This alignment suggests that principal rotational dynamics is preferentially sculpt upon the geometric organization, linking temporal features of the dynamics to anatomical geometry.

## 2 Materials and Methods

### 2.1 Dataset

A dataset of n=20 adult male C57BI6/J mice were previously acquired at the IIT laboratory (Italy). All in vivo experiments were conducted in accordance with the Italian law (DL 2006/2014, EU 63/2010, Ministero della Sanit`a, Roma) and the recommendations in the Guide for the Care and Use of Laboratory Animals of the National Institutes of Health. Animal research protocols were reviewed and consented by the animal care committee of the Italian Institute of Technology and Italian Ministry of Health. Animal preparation, image data acquisition and image data preprocessing for rsfMRI data have been described in greater detail elsewhere [27]. Briefly, rsfMRI data were acquired on a 7.0-T scanner (Bruker BioSpin, Ettlingen) equipped with BGA-9 gradient set, using a 72 mm birdcage transmit coil, and a four-channel solenoid coil for signal reception. Single-shot BOLD echo planar imaging time series were acquired using an echo planar imaging sequence with the following parameters: repetition time/echo time, 1000/15 ms; flip angle, 30°; matrix, 100×100; field of view, 2.3 × 2.3 cm^2^; 18 coronal slices; slice thickness, 0.60 mm; 1920 volumes.

Regarding image preprocessing as described in [28], timeseries were despiked, motion corrected, skull stripped and spatially registered to an in-house EPI-based mouse brain template. Denoising and motion correction strategies involved the regression of mean ventricular signal plus 6 motion parameters [55]. The resulting timeseries were then band-pass filtered (0.01-0.1 Hz band). After preprocessing, mean regional time-series were extracted for 74 (37+37) regions of interest (ROIs) derived from a predefined anatomical parcellation of the Allen Brain Institute (ABI), [46, 65].

### 2.2 Theoretical background

#### 2.2.1 State-space linear model

We considered the standard linear time-invariant continuous stochastic system to model the state dynamics of brain data

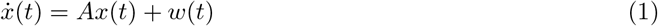

where *x* ∈ ℝ^*n*×1^ is the state vector, *A* ∈ ℝ^*n*×*n*^ is the state interaction matrix, and *w* ∈ ℝ^*n*×1^ is a zero-mean white noise vector with positive definite covariance matrix Σ_*w*_ = 𝔼 [*w*(*t*)*w*(*t*)^⊤^]. In the context of brain dynamics, *A* is generally referred to as the effective connectivity (EC) matrix if the inference is constrained to be physiologically interpretable and plausible. We assumed constant and uncorrelated noise between nodes so that Σ_*w*_ is a scalar matrix of the form Σ_*w*_ = *σ*^2^*I*_*n*_ where *σ*^2^ represents the variance of each node endogenous fluctuation and *I*_*n*_ the identity matrix of size *n*.

Moreover, *A* is assumed to be stable (its eigenvalues have strictly negative real part). This ensures the existence of a positive definite steady-state covariance matrix Σ, which, after normalization, represents the functional connectivity matrix (FC)

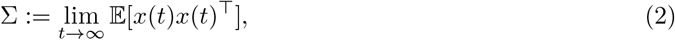

for which the algebraic Lyapunov equation holds

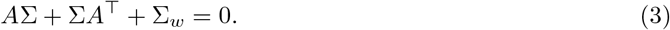

Importantly, *A* can be decomposed such that

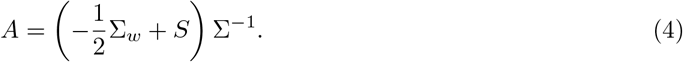

Eq. 4 parametrizes all *A* matrices that give the same Σ, i.e. the covariance matrix of the state, for any *S* skew-symmetric matrix, i.e. *S* = −*S*^⊤^. This defines a one-to-one map between each (Σ, *S*) and *A* given the noise covariance Σ_*w*_ [9], with 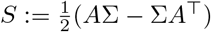.

Considering Eq. 4, we refer to the matrix *A*Σ as the differential covariance [37]. In particular, it holds that 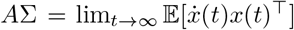, providing a statistical interpretation of the differential covariance. Within our framework, where Σ_*w*_ is diagonal, the two terms −Σ_*w*_*/*2 and *S* decompose the differential covariance into the sum of a diagonal matrix and a hollow matrix (i.e. a matrix with zeros on the main diagonal). We refer to the former as the differential auto-covariance (dA-Cov), and to the latter as the differential cross-covariance (dC-Cov). As a consequence, each off-diagonal element *S*_*i,j*_ admits the statistical interpretation 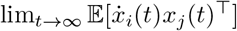, and these terms account entirely for the asymmetric component of *A*. Indeed, the following algebraic Lyapunov equation also holds

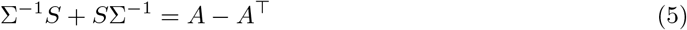

Note that if correlated noise is present, as not considered in this study, dC-Cov and *S* are no longer interchangeable.

On the interpretation of dC-Cov and dA-Cov, we refer to our previous work [5]. Here, we remark that a non-zero *S* matrix introduces time irreversibility into the dynamics. Moreover, inverting the sign of *S* corresponds to reversing the direction of the dynamics [12]. This provides a direct interpretation of the system in terms of source and sink nodes, identifying which nodes act as drivers and which as recipients of the dynamical flow [37, 10].

#### 2.2.2 Lagged covariance and power spectral density

The matrix *S*, encoding endogenous time-lag interactions, directly determines the structure of the lagged covariance matrix Σ(*τ*) := lim_*t*→∞_ 𝔼 [*x*(*t* + *τ*)*x*(*t*)^⊤^] and its Fourier transform, i.e. the power spectral density matrix Φ(*ıω*). In particular, a non-zero *S* matrix gives rise to asymmetries in the time-lagged covariance matrix and Hermitian symmetry on the complex cross-spectral density matrix. Both quantities can be written in a closed form when dealing with a linear state-space model:

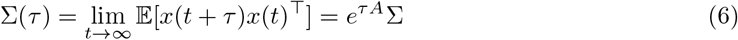

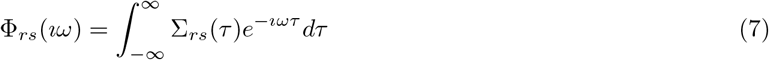

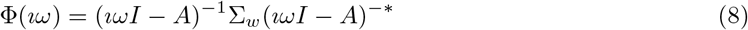

where ^*^ stands for the conjugate transpose, and *ı* for the imaginary unit. If *S* = 0, the differential covariance matrix satisfies the symmetry condition *A*Σ = Σ*A*^⊤^. By induction, this relation extends to any power of *A*, i.e. *A*^*k*^Σ = Σ(*A*^*k*^)^⊤^

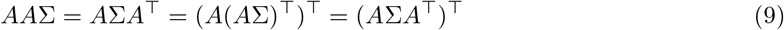

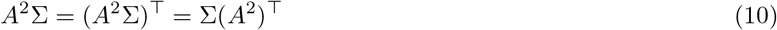

Since the matrix exponential admits the power series expansion 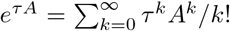, it follows that *e*^*τA*^Σ = Σ(*e*^*τA*^)^⊤^ for any lag *τ*. In other words, in the absence of *S*-driven interactions, the lagged covariance matrix remains symmetric at all time scales. Moreover, given that Σ_*rs*_(*τ*) = Σ_*sr*_(−*τ*), it follows that if the lagged covariance matrix is symmetric, then Σ_*rs*_(*τ*) is an even function of the lag *τ*. As a consequence, its Fourier transform Φ_*rs*_(*ıω*) which represents the cross-spectral density, is a real and even function of frequency. This observation highlights the role of the asymmetric component of 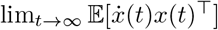, encoded by the *S* matrix. Specifically, the presence of a non-zero *S* breaks the time-reversal symmetry of the dynamics, inducing asymmetries in the lagged covariance (Eq. 6), and generating a complex-valued power spectral density (Eq. 7). Note that this also holds in the presence of a correlated noise covariance Σ_*w*_. However, it is important to emphasize that *S*_*rs*_ = 0 does not necessarily imply that the corresponding Σ(*τ*) and Φ(*ıω*) entries are even and real, respectively.

An alternative perspective to interpret the differential covariance is

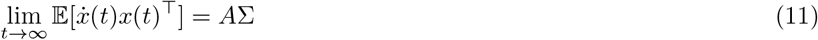

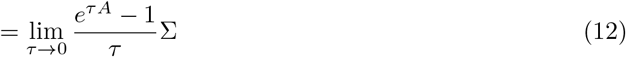

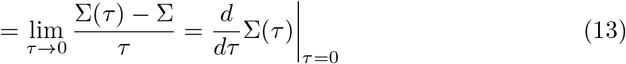

so as to obtain that *A*Σ is the derivative of Σ(*τ*) evaluated at *τ* = 0. Symmetric and nonzero entries in *A*Σ imply that the derivative of Σ(*τ*) with respect to *τ* has an opposite sign when approaching *τ* = 0 from the positive and negative sides. However, in our modeling framework, symmetric entries of *A*Σ correspond only to zero entries of *S*. As a result, differentiability at *τ* = 0 is preserved, making zero lag a local maximum or minimum of the lag covariance. In contrast, skew-symmetric entries in *A*Σ imply that the derivative of Σ(*τ*) with respect to *τ* remains constant and nonzero at *τ* = 0, resulting in a smooth evolution of the lag covariance without a local extremum at zero lag.

#### 2.2.3 Differential cross-covariance decomposition

To investigate the structure of the lagged interactions encoded in the matrix *S*, the Schur decomposition offers a powerful analytical tool. In particular, it enables the identification of rotational modes that share the same angular frequency, thus revealing coherent dynamical patterns within the system. When applied to a real square matrix *R* ∈ ℝ^*n*×*n*^, it yields

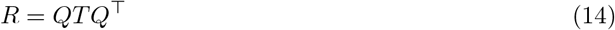

where *Q* ∈ ℝ^*n*×*n*^ is an orthogonal matrix, and *T* ∈ ℝ^*n*×*n*^ is an upper quasi-triangular matrix, i.e. a block upper triangular matrix with either real entries on the diagonal for real eigenvalues, or [2 × 2] blocks corresponding to complex conjugate eigenvalue pairs. In the special case where *R* is a real skew-symmetric matrix, all eigenvalues are purely imaginary or zero, and the Schur form reduces to a block-diagonal matrix composed of real [2 × 2] skew-symmetric blocks of the form:

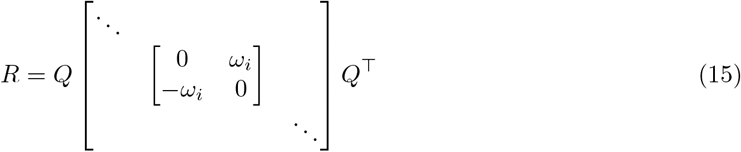

where *λ*_*i*_ = ±*ıω*_*i*_ are the purely imaginary complex conjugate eigenvalue pairs. Moreover, recalling that the matrix exponential of each [2 × 2] block is:

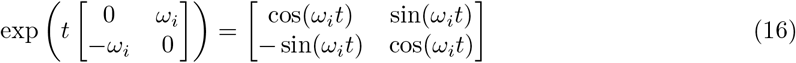

this decomposition highlights the presence of rotational components in the dynamics encoded by the dC-Cov matrix. Each [2 × 2] block captures a mode rotating in a 2D plane with angular frequency *ω*_*i*_, generating a clockwise rotation when *ω*_*i*_ *>* 0. Given an initial condition *x*_0_, the circular trajectories defined by *S* are *x*(*t*) = exp(*tS*)*x*_0_. Since this solves the linear differential equation 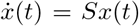 it results that this latter is the velocity field at any point *x*(*t*). A drawback of this decomposition is its non-uniqueness. Indeed, given a Schur mode defined by a pair of orthogonal vectors [*q*_1_ *q*_2_] ∈ ℝ^*n*×2^and an angular frequency *ω*, the following holds

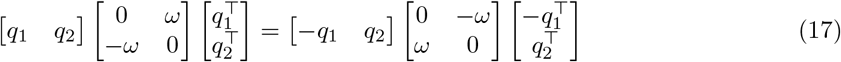

which implies that the decomposition does not uniquely specify which of the two orthogonal vectors acts as the sender and which as the receiver.

#### 2.2.4 Spatiotemporal lagged interactions

The complex spatiotemporal pattern of lagged interactions arises through the product *S*Σ^−1^, which introduces a warping of the velocity field. The axes along which this deformation occurs correspond to the eigenvectors of the precision matrix Σ^−1^. Considering Σ^−1^ = *V*Δ*V*^⊤^ with *V* and Δ its eigenvectors and eigenvalues, respectively, we get

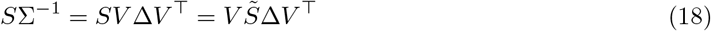

where 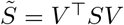 is still skew-symmetric. From Eq. 5 and given *P* = *V* ^⊤^(*A* − *A*^⊤^)*V*, we get

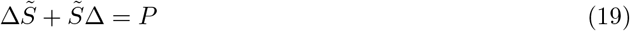

which being 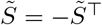 and Δ diagonal, it can be written as

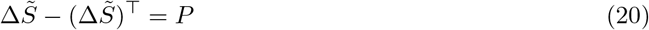

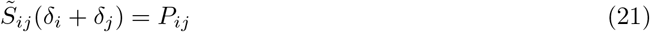

where *δ*_*i*_ is the *i*-th eigenvalue in Δ, Eq. 21 gives an entry-wise formula that relates the asymmetric part of *A* and matrix *S* in the eigenbasis of Σ^−1^.

However, by projecting on the eigenvectors of the precision matrix does not in general allow acting on the single rotational model independently. Consider the Schur decomposition of *S* instead, and for simplicity assume that the precision matrix on the Schur base shows the same block-diagonal structure of the matrix *S*. This decomposes the dynamics as an independent sum of 2D rotational modes. Focus on a 2-dimensional system with

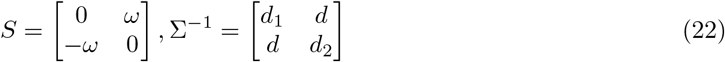

where *ω >* 0 is the angular speed of a clockwise rotation, and *d*_1_ *>* 0, *d*_2_ *>* 0, *d*^2^ *< d*_1_*d*_2_ from the positive definiteness of Σ^−1^. Consider they product

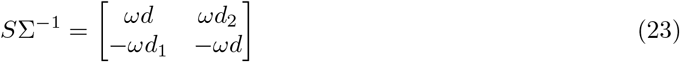

by computing the roots of the related characteristic polynomial *p*(*λ*) = det(*λI* − *S*Σ^−1^), we obtain the eigenvalues 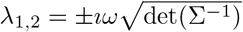. Thus, Σ^−1^ scales the angular speed of the rotation proportionally to its determinant. The scaling along the two new basis of the Schur modes of *S*Σ^−1^ depends on the eigenvalues of Σ^−1^, if they differ the rotation becomes elliptical. Also the phase of the two trajectories can be altered by Σ^−1^. Initially they are in phase quadrature, i.e. Δ*ϕ* = *π/*2 being *S* skew-symmetric, and how the phase diverge from quadrature depends on the eigenvectors of *S*Σ^−1^. Consider the eigenvalue 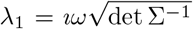, and its related eigenvector *v*_1_ = [*v*_11_, *v*_21_]^⊤^, the phase shift is given by the argument of the ratio *v*_11_*/v*_21_:

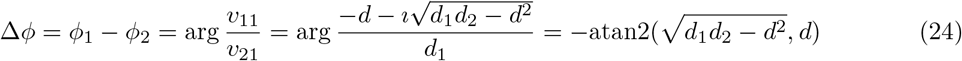

having defined in Eq. 22 the matrix *S* as a clockwise rotation, the phase shift must be interpreted as a delay of *x*_1_, thus *x*_1_ lags *x*_2_ by Δ*ϕ* radians. In particular, a phase shift occurs only if *d* ≠ 0, i.e. Σ^−1^ is not diagonal.

In the general case, where the transformation of Σ^−1^ into the Schur basis of *S* does not yield a [2 × 2] block-diagonal matrix, the effect of the precision matrix is to couple different Schur modes of *S*. As a result, new rotational Schur modes emerge. Indeed, since *S*Σ^−1^ is similar to Σ^−1*/*2^*S*Σ^−1*/*2^, and the latter is skew-symmetric with eigenvalues ±*ıµ*_*k*_, the matrix *S*Σ^−1^ has only purely imaginary eigenvalues.

So far, we have only considered rotational modes by focusing on the product *S*Σ^−1^. To introduce a dissipative component, we consider the matrix *ξI* + *S*. Let *V* and Ω denote the Schur decomposition of *S*, such that *SV* = *V* Ω. It follows that (*ξI* + *S*)*V* = *V* (*ξI* + Ω). Therefore, the Schur orthogonal vectors are preserved, while the eigenvalues are shifted to *ξ* ± *ıω*_*i*_. From Eq. 4, we have *ξ* = −*σ*^2^*/*2. Its physical interpretation is a decay rate that defines the timescale of the rotational modes. Initially, this decay is assumed to be the same across all modes, while the multiplication with Σ^−1^ introduces heterogeneity in the decay rates.

### 2.3 sparse-DCM

The effective connectivity matrix, along with other model variables, was estimated at the single-subject level using the method described in [50], which is referred to as sparse-DCM. In line with the DCM framework [21], sparse-DCM is a state-space model where the state *x*(*t*) satisfies a set of linear differential equations representing the coupling among neural components, and the output model maps the neuronal activity to the measured BOLD signal *y*(*t*) through the hemodynamic response function (HRF):

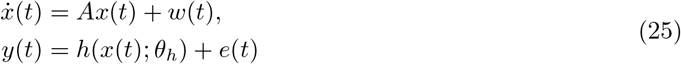

with *A* representing the effective connectivity matrix, *h*(.) the hemodynamic response that is modeled by the biophysically inspired Balloon-Windkessel model [22] and *θ*_*h*_ its parameters. *w*(*t*) denotes the stochastic intrinsic brain fluctuations and *e*(*t*) the observation noise, both are Gaussian variables with zero mean and diagonal covariance matrices *σ*^2^*I*_*n*_ and *R* = diag(*λ*_1_, *λ*_2_, …, *λ*_*n*_), respectively.

To address the computational burden of model inversion when dealing with whole brain data, in [50] a discretization and linearization of Eq. 25 as well as a sparsity-inducing prion on *A* was proposed. This was motivated by the low temporal resolution of fMRI data, which usually ranges from 0.5 to 3 seconds, and the idea that the hemodynamic response *h*(·.) can be modeled as a Finite Impulse Response (FIR) model with input the neuronal state and output the BOLD signal. In our study, to ensure that the length of the input response was large enough to model relevant temporal dependencies, we set the hemodynamic length to 18 samples with a sampling time 1s (TR). For each brain parcel *i*, a finite impulse response *h*_*i*_ ∼ 𝒩 (*µ*_*h*_, Σ_*h*_) was assigned by deriving *µ*_*h*_ and Σ_*h*_ through a Monte-Carlo sampling of typical responses generated by the non-linear Balloon-Windkessel model (10000 samples). The sparsity-inducing prior on the EC estimation was formulated to reduce as much as possible spurious couplings. In particular, each element *a*_*i*_ of matrix *A* was assumed to be a Gaussian variable with zero mean and *γ*_*i*_ variance. The hyperparameter 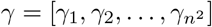 was estimated through marginal likelihood maximization. Under generic conditions, the maximum likelihood estimate of some *γ*_*i*_-s will be zero such that the Gaussian posterior distribution of their corresponding *a*_*i*_ is concentrated around zero thus producing a zero MAP estimate. In sparse-DCM, model inversion and parameter optimization are performed by an expectation-maximization (EM) algorithm.

### 2.4 Metric description

#### 2.4.1 Distance matrix and Laplacian spectral decomposition

The distance between each pair of regions was computed as the Euclidean distance between region centroids. Each centroid was defined as the mean of the *x, y* and *z* coordinates of all voxels belonging to the region. The resulting distance matrix was then converted into a proximity matrix using a Gaussian (rbf) kernel with *σ* = 500. In our case, this choice ensured that the shortest 20% of distances corresponded to proximity values above 0.5, i.e. within the upper half of the kernel range. We refer to that percentage (20% in the example above) as the target density. We then decomposed the proximity matrix derived from spatial distances using the Laplacian spectral decomposition technique. Specifically, we adopted the degree-normalized Laplacian,

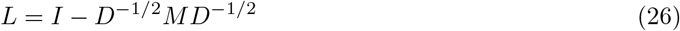

where *M* denotes the proximity matrix and *D* the diagonal degree matrix of *M*. The spectral decomposition of *L*, i.e. *L* = *U* Π*U* ^⊤^, provides the orthogonal basis *U*, whose column vectors correspond to the Laplacian harmonics of *M*.

In the context of network dynamics, the degree-normalized Laplacian *L* can be related to the random-walk Laplacian *L*_*rw*_ [36]. In particular, the following similarity transformation holds: *L*_*rw*_ = *D*^−1*/*2^*LD*^1*/*2^, which implies that the two operators share the same spectrum. *L*_*rw*_ admits a natural interpretation in terms of weighted consensus dynamics: 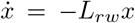 where, for each node, 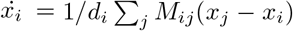. This equation shows that each node evolves toward a weighted average of its neighbors’ states, with interaction strengths normalized by its degree. By applying the change of coordinates *y* = *D*^1*/*2^*x* the dynamics can be written as 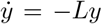 which corresponds to a consensus process governed by the degree-normalized Laplacian, i.e. in a degree-weighted coordinate system. As a consequence, the final consensus state is not uniform across nodes but is instead weighted by *D*^1*/*2^, reflecting the network’s degree heterogeneity (see Figure 3d).

From a complementary perspective, the degree-normalized Laplacian can be interpreted through the lens of spectral graph theory, where it is closely related to the notion of normalized cut (Ncut) and graph bi-partitioning [4]. In this setting, given a graph *G* = (*V, E*) with adjacency matrix *M*, consider a bi-partition of the node set into two disjoint subsets *A* and *B* such that *A* ∪ *B* = *V* and *A* ∩ *B* = ∅. The cut between the two sets measures their dissimilarity by summing the total weight of edges that connect nodes across the partition:

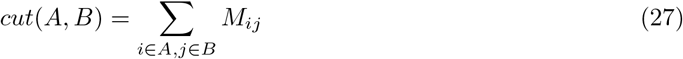

To account for partition unbalancing, the normalized cut [60] was proposed:

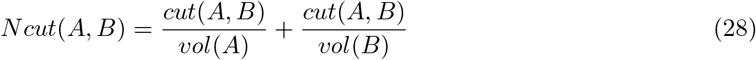

where *vol*(*S*) =Σ _*i*∈*S*_ *d*_*i*_ is the total connectivity of the nodes in *S*, and *d*_*i*_ denotes the degree of node *i* (i.e. the *i*-th diagonal entry of the degree matrix *D*). This combinatorial formulation can be expressed in algebraic form by introducing an indicator vector *f* that encodes the partition: 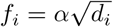 or 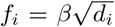 if *i* ∈ *B* with 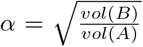 and 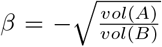. The normalized cut can be written in compact form:

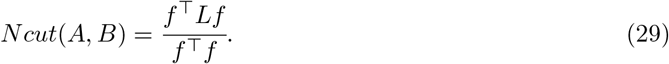

The problem of optimal bi-partitioning the graph encoded by *M* can be formulated as the minimization of the normalized cut. However, this problem is known to be NP-complete. A common approximation is obtained by relaxing the discrete bi-partition to a continuous indicator vector *x*, and imposing *x* ⊥ *D*^1*/*2^**1**. For the degree-normalized Laplacian, it holds:

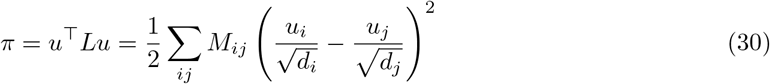

for any eigen-pair (*u, π*) of *L*. Since *L* is positive semidefinite, the eigenvector associated with the second smallest eigenvalue minimizes Eq. 30 among all vectors orthogonal to *D*^1*/*2^**1**. Consequently, this eigenvector provides a natural continuous relaxation of the normalized cut problem, and can be used to approximate a minimizer of Eq. 29. To recover a discrete partition from the relaxed solution, the simplest way is to threshold the eigenvector by its sign: nodes with *u*_*i*_ *>* 0 are assigned to set *A*, while nodes with *u*_*i*_ *<* 0 are assigned to set *B*. Higher-order eigenvectors can be used in a similar manner to defined additional partitions; however, these solutions are constrained to lie within the orthogonal eigenvector basis of *L*. As a result, moving to higher eigenvectors explores progressively finer partitions of the graph, but without guaranteeing optimality with respect to the normalized cut objective.

#### 2.4.2 Participation ratio

The participation ratio (pr) is a positive index that quantifies the degree of localization of a mode, as reflected in its eigenvector [11]. Let *v* ∈ ℝ^*n*×1^ denote the eigenvector associated with a given mode, its participation rate is defined as

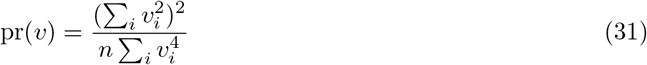

which reaches its maximum value of 1 when the mode is minimally localized, i.e. uniformly distributed across all nodes. For normalized eigenvectors, where *v*^⊤^*v* = 1, this simplifies to

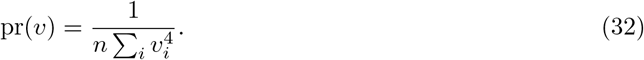

#### 2.4.3 Harmonic power

Given an eigenvector of the Laplacian matrix *v* ∈ ℝ^*n*×1^, each element of its harmonic power 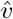 was defined as in [62]

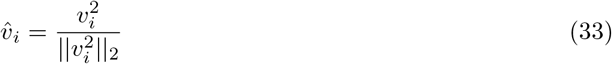

#### 2.4.4 Coherency and coherence

Given the cross-spectral density matrix as in Eq. 8 and following the terminology in [44], the coherency between the pair (*i, j*) of a given angular frequency *ω* is the normalized cross-spectrum

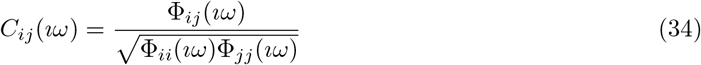

and it quantifies the phase coupling between the two signals, the coherence is its absolute value

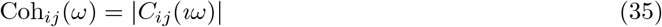

## 3 Results

Results are organized into two sections: the first focuses on simulations designed to illustrate the roles of the symmetric and anti-symmetric components of the differential covariance, while the second examines the relationship between inter-region distances and the corresponding elements of the dC-Cov matrix.

### 3.1 Illustrative simulations

Panels in Figure 1 show three examples of three dimensional state-space simulations with predefined covariance Σ and dC-Cov *S*. In all cases, the noise covariance is diagonal and of the form Σ_*w*_ = *σ*^2^*I*_*n*_.

**Figure 1:**
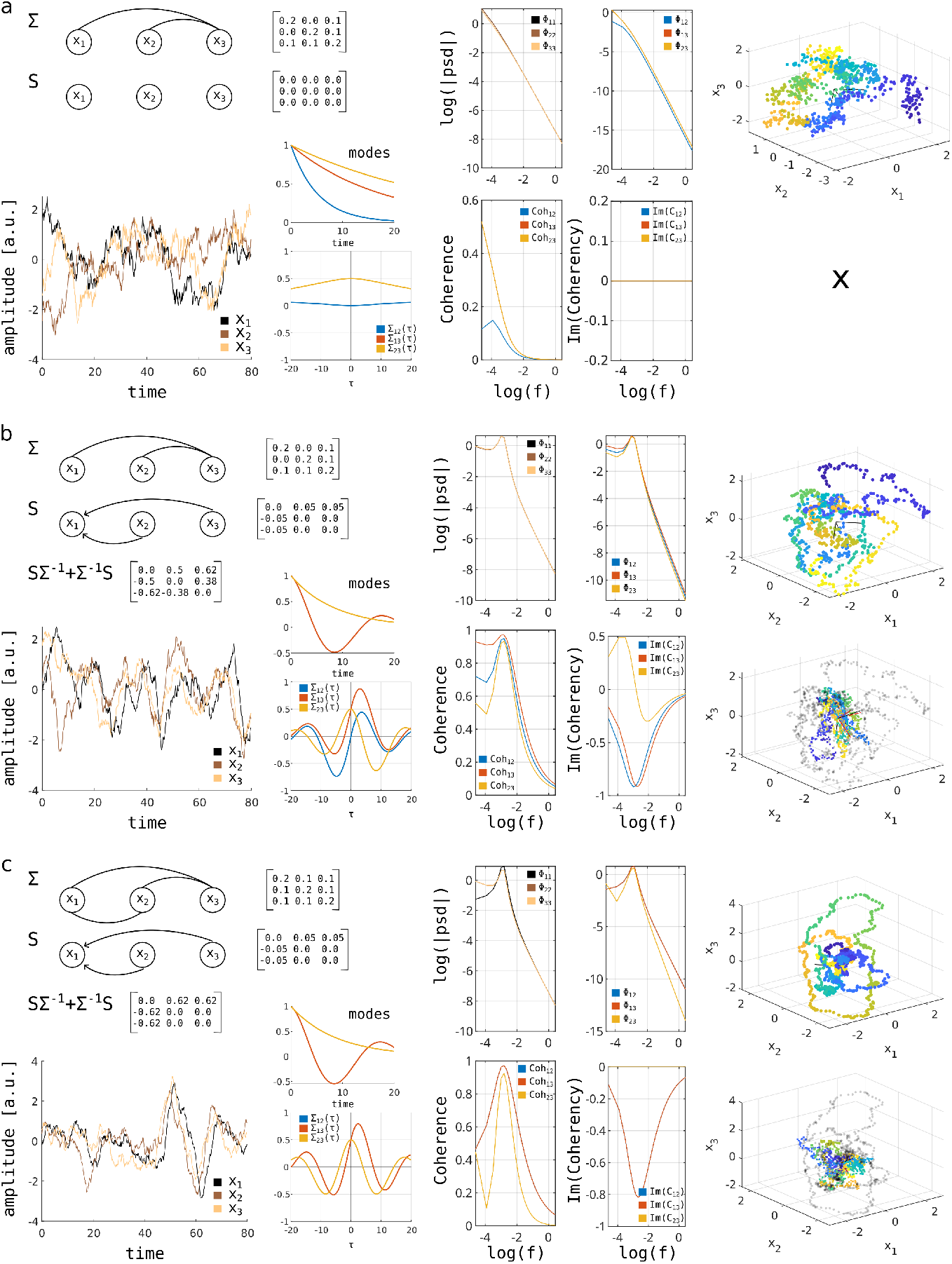
Explanatory panels highlighting how the differential covariance and covariance matrices contribute to determining the dynamics of a 3-dimensional linear system. Each panel reports: the corresponding Σ and *S* matrices, and the resulting modes of the system Jacobian; an example of a signal realization with the corresponding lag-covariances Σ_*ij*_(*τ*) for each pair *ij*; and, in the frequency domain, the log-magnitude of the power spectral density (both auto- and cross-), along with the cross-coherence and the imaginary part of coherency. On the right, simulated signals are shown both in the original data space and projected onto the corresponding pair of Schur basis vectors. These results are shown under three simulation assumptions: panel *a*, a time-reversible system (*S* = 0); panel *b*, a general time-irreversible system; panel *c*, a special case of a time-irreversible system with no distortion of the rotational plane.

Panel *a* assumes no anti-symmetric component, *S* = 0, meaning that the system operates at thermodynamic equilibrium. Evidences of this is provided by the following observations: (i) all modes decay exponentially, (ii) the lag-covariances Σ_*ij*_(*τ*) are even functions, (iii) the power spectral densities Φ_*ij*_(*ıω*), both within and across signals, are monotonically decreasing functions, and (iv) the imaginary part of the coherency Im(*C*_*ij*_) is zero at all frequencies, indicating a phase difference of zero between signals across the spectrum.

In panel *b*, we added time-lagged interactions from signal 3 to 1 and from signal 2 to 1. This modification is evident in the system modes, as one now behaves as a damped oscillator. The lag-covariances are not even functions, reflecting the presence of non-zero time lags that maximize the signal covariance. In the frequency domain, this results in cross-spectrum peaks at non-zero frequencies, with the imaginary part of the coherency capturing the corresponding phase lags. An additional point of interest is how the time-lagged interactions induced by *S* affect the pair (2, 3), which is not directly connected. Although its lag-covariance Σ_23_(*τ*) has a zero derivative at *τ* = 0 (imposed by *S*_23_ = *S*_32_ = 0), it is still not an even function due to the distortion caused by the covariance matrix. This implies a complex cross-spectrum for this pair, with Im(*C*_23_) ≠ 0. Finally, note also that even though *S*_12_ = *S*_13_, the peaks of their lag-covariances are not aligned at the same *τ* ^*^, and their corresponding Im(*C*) components peak at different frequencies. This is due to how the precision matrix Σ^−1^ alters the rotational modes encoded in the matrix *S*. In this example, the rotational plane is tilted, causing a mixing between modes. To illustrate this, the panel also shows the antisymmetric part of the product *S*Σ^−1^ (which according to our modeling assumption corresponds to the antisymmetric part of *A*), highlighting that the skew structure of *S* is no longer preserved. Equivalently, *S*Σ^−1^ does not block-diagonalize on the Schur basis of *S*. On the right side of the panel, scatter plots of the three simulated signals are shown, both in the original data space and projected onto the corresponding pair of Schur basis vectors.

The last panel preserves the same matrix *S* but modifies the covariance Σ to counteract the effects observed in the previous panel. As a result, all pairwise lag-covariance functions become even, and Σ_12_(*τ*) = Σ_13_(*τ*). In terms of Im(*C*), the pair (2, 3) is zero at all frequencies, while the other two overlap. The chosen Σ^−1^ preserves the rotation plane of *S*, only rescaling the rotation speed. As a result, the Schur basis of *S* remains a block-diagonalizing basis for *S*Σ^−1^, and all signal pairs undergo the same phase shift.

To better highlight the role of the precision matrix and in particular how it deforms the rotational matrix *S* through the product *S*Σ^−1^, we focused on a 2-dimensional example, see Figure 2. This is a numerical example of what is described in Section 2.2.4. The figure is composed of 3 panels, in all cases the matrix *S* is defined to reproduce a clockwise rotation of angular speed *ω* = 2*π*. In panel *a*, the precision matrix is the identity matrix so it does not deform the original circular rotation. As a consequence, the two signals have a phase difference Δ*ϕ* = *π/*2, i.e. a time lag of 0.25*s* being *f* = 1Hz.

**Figure 2:**
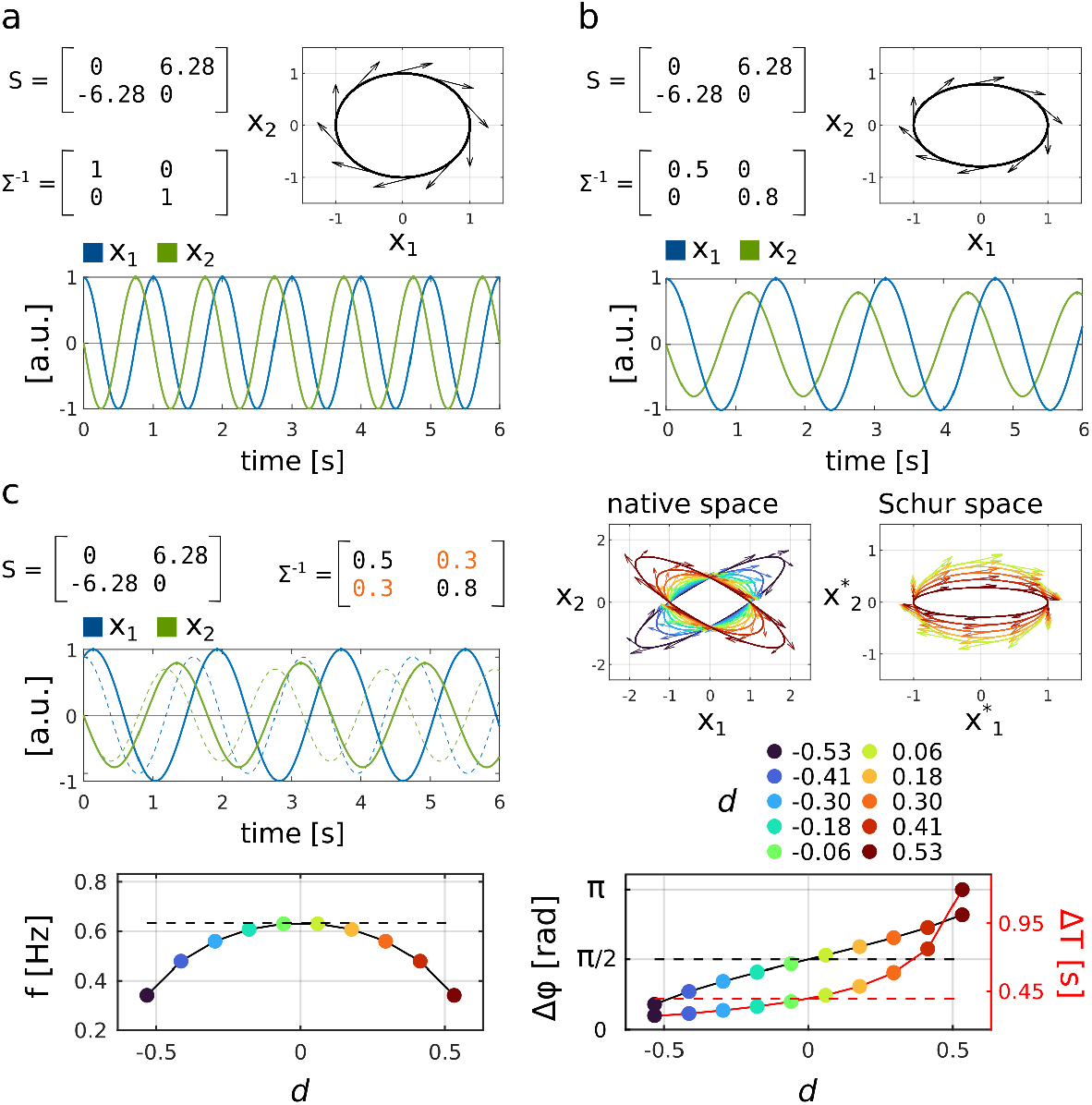
Deformation of the same two-dimensional rotational model induced by multiplication with the precision matrix Σ^−1^. Panel *a*: absence of deformation, corresponding to a circular mode with exact quadrature. Panel *b*: transition from circular to elliptical rotation while preserving quadrature, this modifies the angular frequency and consequently the associated time lag. Panel *c*: the introduction of a non-zero off-diagonal entry *d* in Σ^−1^ modulates both the angular frequency and the phase lag.

Panel *b* introduces a change in the diagonal entries of Σ^−1^, (*d*_1_, *d*_2_). The effect is to independently alter the velocity field along the two dimensions. The circular trajectory becomes elliptical, and the overall angular speed scales with the geometric mean of the diagonal entries, 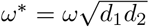. Note that the phase in radian is preserved, but changing the frequency changes the time lag as well.

Panel *c* includes non-zero off-diagonal entry *d* in the precision matrix. To preserve the positive definiteness of Σ^−1^ the condition *d*^2^ *< d*_1_*d*_2_ must hold. A full precision matrix modifies the Schur form of *S*, altering both frequency and phase lag. The panel shows the specific case of *d* = 0.3, as well as the effects of varying *d* within its admissible bounds. The frequency decreases as the absolute value of the off-diagonal entry increases (the dashed horizontal line marks the maximal frequency obtained at *d* = 0). The phase lag increases with *d*, approaching 0 at the minimum admissible value of *d*, and *π* as *d* approaches its upper bound. The time lag Δ*T* shows a similar trend, although the relationship is not linear.

### 3.2 Relating spatial distance to single-subject Schur modes

The second part of the Results section focuses on empirical data, examining the relationship between 2 × 2 Schur modes and spatial properties of mouse brain regions. Structural connectivity is often used to constrain brain dynamic, with functional strength between regions weighted proportionally to their structural strength. By contrast, geometrical properties such as the spatial distance between regions are generally less explored. We hypothesize that the Schur modes encoded in the *S* matrix may reflect geometrical properties of the brain, such as inter-regional distances. In line with this assumption, we derived a pairwise distance matrix from the mouse brain after a parcellation in 74 regions. Resting-state fMRI signals from 20 mice were then merged using the same parcellation scheme. Figure 3a shows the 74 parcels and their centroids, which were used to compute the pairwise Euclidean distance matrix. This distance matrix was converted into a proximity matrix using a Gaussian (rbf) kernel; the resulting proximity matrix and kernel are shown in Figure 3b. The bottom-right part of the same panel also reports the eigenvalue spectrum of the normalized Laplacian computed from the proximity matrix, together with the corresponding eigenvector participation ratios. The spectrum of the normalized Laplacian admits a direct interpretation in terms of the normalized cut, which is reported in Figure 3c for each eigenvector. From a complementary dynamical perspective, the normalized Laplacian also defines the consensus dynamics shown in Figure 3d. The same panel displays the first seven leading eigenvectors, highlighting the edges within each partition according to the sign of the eigenvector components.

**Figure 3:**
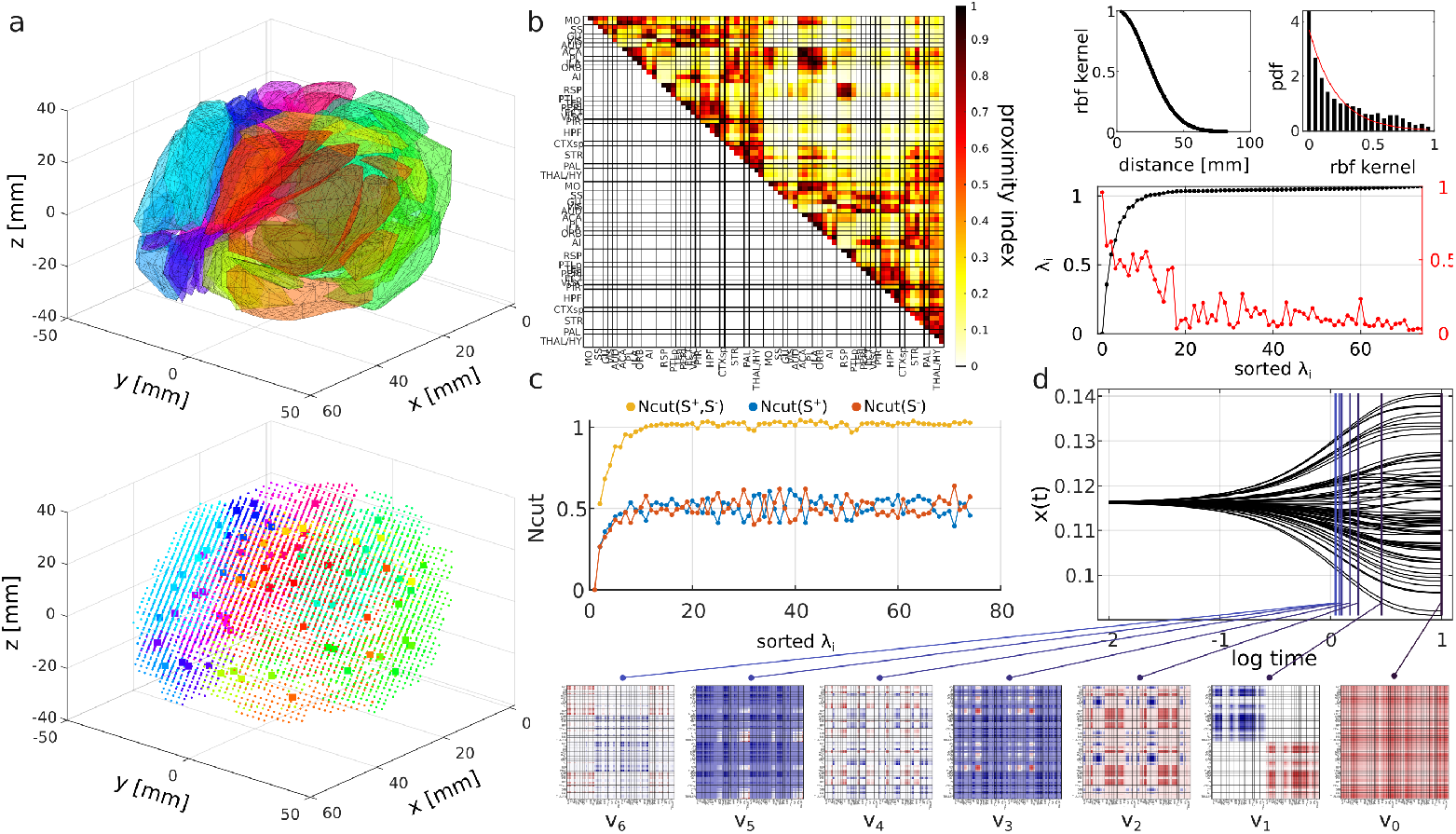
Mouse brain geometry and spectral analysis. Panel *a*: three-dimensional triangulation reconstructed from the *x, y, z* coordinates of each voxel obtained from magnetic resonance imaging (top). The bottom panel shows the same geometry, highlighting individual voxels (small markers) and the centroids of each parcel (large markers) derived from the anatomical atlas used to cluster the full volume; colors indicate the parcellation. Panel *b*: Euclidean distance matrix transformed into a proximity matrix using a Gaussian kernel with predefined *σ*. The corresponding kernel profile and the distribution of proximity values are shown in the top-right plots, while the plot below reports the eigenvalue spectrum together with the associated participation ratio. Panel *c*: normalized cut values computed for each eigenvector (sorted by increasing eigenvalues), where partitions are obtained by thresholding the eigenvector entries according to their sign. Panel *d* : consensus dynamics generated by the normalized Laplacian. The first seven eigenvectors are mapped to partitions: entries sharing the same color correspond to nodes within the same cluster, whereas white entries denote edges between clusters. Cluster separation is determined by the sign of the eigenvector components.

For the resting-state fMRI data, the DCM model described in Section 2.3 was identified separately for each mouse data, and the corresponding *S* matrix was Schur decomposed as in Section 2.2.3. Figure 4a shows the frequency spectrum of the Schur modes across subject. The frequency of the faster modes displayed large inter-subject variability, whereas beyond the first few modes it decreased rapidly and became consistent across subjects. We next examined the variability of the Schur-mode eigenvectors across subjects. Focusing on the first two high-frequency modes, we found that their optimal matching for each subject pair showed a lower mean index difference compared with that obtained from randomly selected modes. For any two subjects, the index difference between a pair of optimally matched eigenvectors refers to the gap in their positions after sorting the eigenvectors by increasing eigenvalues. The matching procedure was based on an objective function that minimized the orthogonality between corresponding eigenvectors across subjects. This effect is visible in the upper triangular matrix of Figure 4b, whereas randomly selected paired modes (lower triangular matrix) displayed higher mean index gap. A complementary analysis using inner products (the score of the objective function) between optimally matched eigenvectors showed that the highest-frequency modes not only align in index but also exhibit the largest inner product scores compared to modes randomly drawn from across the full frequency spectrum, Figure 4c.

**Figure 4:**
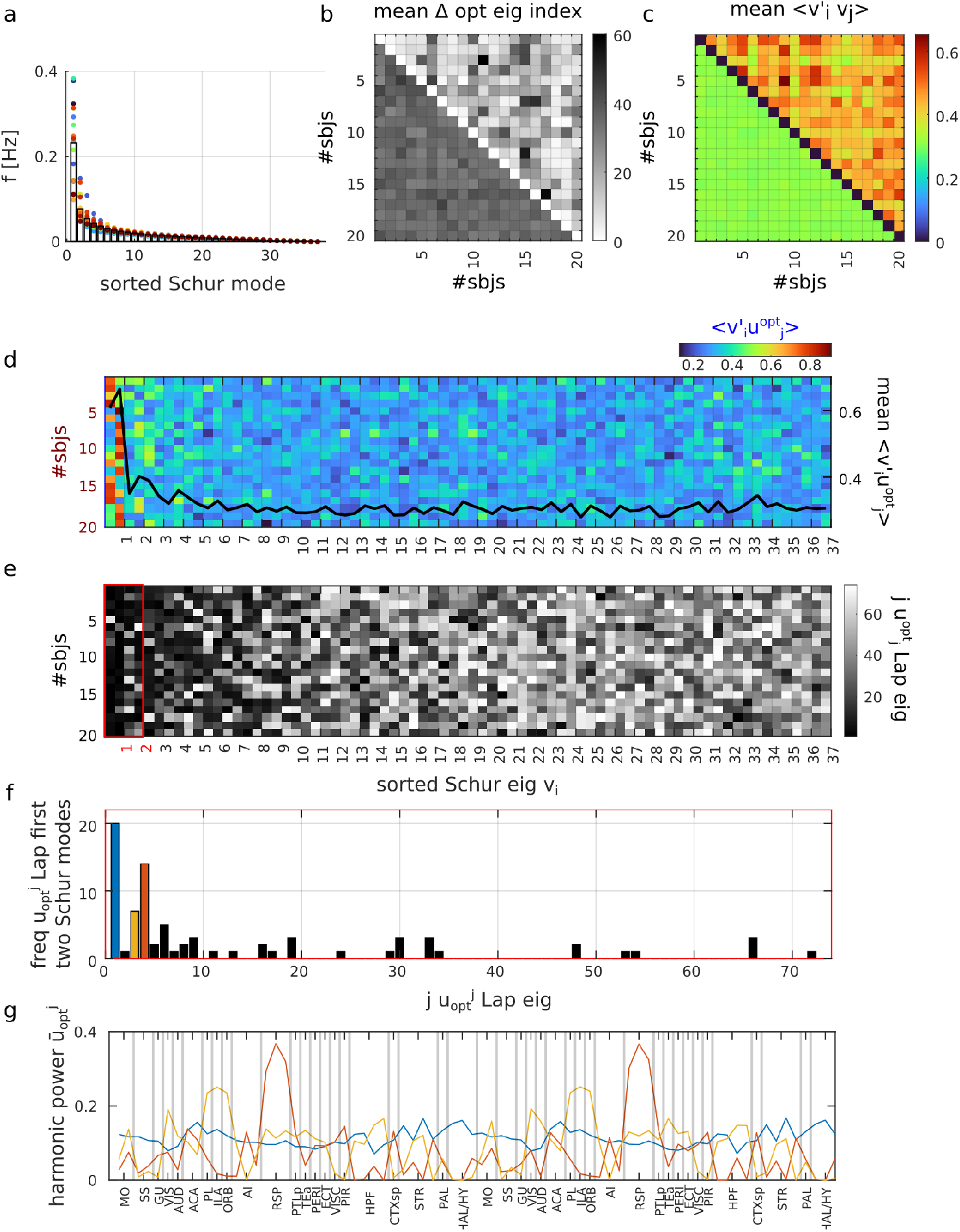
Alignment between normalized Laplacian eigenvectors and single-subject Schur modes. Panel *a*: sorted angular frequencies of the Schur modes for each mice (color coded). Panel *b*: for each mouse pair, the mean index difference between optimally matched Schur vectors (Δ opt eig index) within the two dominant Schur modes (upper triangular part), compared with that obtained from two randomly selected pairs (lower triangular part; average over 100 sampling repetitions). Panel *c*: same as panel *b*, but reporting the inner product between optimally matched Schur vectors 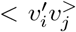. The upper triangular part shows the mean value over the two dominant Schur modes, while the lower triangular part reports the mean over randomly selected pairs (average over 100 repetitions). Panel *d* : coupling measure defined as the inner product between the optimally matched Laplacian eigenvectors 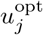 and the Schur vectors *v*_*i*_ (ordered by decreasing angular frequencies) for each mouse. Panel *e*: index *j* ∈ ℤ| 1 ≤ *j* ≤ 74 (black-white colormap), of the optimally matched Laplacian eigenvector 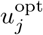 associated with the *i*-th Schur vectors (x-axis). Panel *f* : number of occurrences across mice for each Laplacian eigenvector within the first two dominant Schur modes (red box in panel *e*). Panel *g* : harmonic power vectors of the three most frequently assigned Laplacian eigenvectors.

Building on the idea of optimal alignment between two sets of eigenvectors, we quantified the coupling between Laplacian eigenvector of the spatial distance matrix and each subject’s Schur modes. Figure 4d reports, for all subjects (rows), the inner product between each Schur mode (columns) and its optimally paired Laplacian eigenvector. Schur modes are sorted in descending order of frequency. On the left y-axis, we show the across-subject average of the inner product values for each Schur mode. The strongest couplings were consistently found in the higher-frequency modes. Figure 4e shows the index of the Laplacian eigenvector (previously sorted by eigenvalue) assigned to each Schur mode by the linear assignment solver. This analysis revealed that faster Schur modes were generally paired with the most dominant Laplacian eigenvectors, i.e. those associated with lower eigenvalues. To better understand what these most frequently assigned Laplacian eigenvectors encode, we analyzed their occurrence and spatial patterns. Figure 4f reports, for each Laplacian eigenvectors, the frequency with which it was assigned to the first two Schur modes (four Schur vectors in total) across subjects. The three most frequently assigned eigenvectors are illustrated in Figure 4g through their harmonic power vectors. The first Laplacian eigenvector, which had the highest number of matches, corresponds by definition to the degree of the spatial distance matrix. The second most frequent match was the 4th Laplacian eigenvector, which primarily spans the retrosplenial area. The third corresponds to an eigenvector encompassing the prelimbic, infralimbic and orbital areas. Notably, these regions form the midline default mode network.

We then asked whether the structural connectivity matrix would exhibit a similar alignment with the Schur basis vectors. To address this question, we repeated the same analysis by applying the normalized Laplacian transformation to the symmetric mouse structural connectivity across different target densities, achieved by varying the variance of the Gaussian kernel (see Section 2.4). Figure 5a (top panels) reports, for each mouse, the mean coupling between the dominant Schur modes (top two mode pairs) and their optimally matched Laplacian eigenvectors as a function of the target density. In the left panel, the mean matching score obtained using the distance matrix is shown in red, while the corresponding score obtained using the structural connectivity matrix is shown in blue. The right panel reports the relative difference between distance-based and structure-based matching scores. At low target density (i.e. high sparsity level), the two matching are comparable. However, as density increases (i.e. sparsity is reduced), the distance matrix exhibits a stronger alignment with the Schur modes, with statistically significant differences observed at 30%, 40% and 50%. These percentages correspond to the fraction of shortest-distance entries (or, equivalently, strongest structural connections) mapped into the upper half of the kernel range. Similarly, we repeated this analysis by matching both the distance-based similarity and the structural connectivity matrix with the subject-specific covariance matrix, i.e. the unnormalized functional connectivity. This served as a sanity check to confirm the expected stronger alignment between covariance with structural connectivity. The corresponding results are shown in Figure 5a (bottom panels). As in the previous analysis, the difference increased with the target density, reaching statistical significance at the four highest density levels.

**Figure 5:**
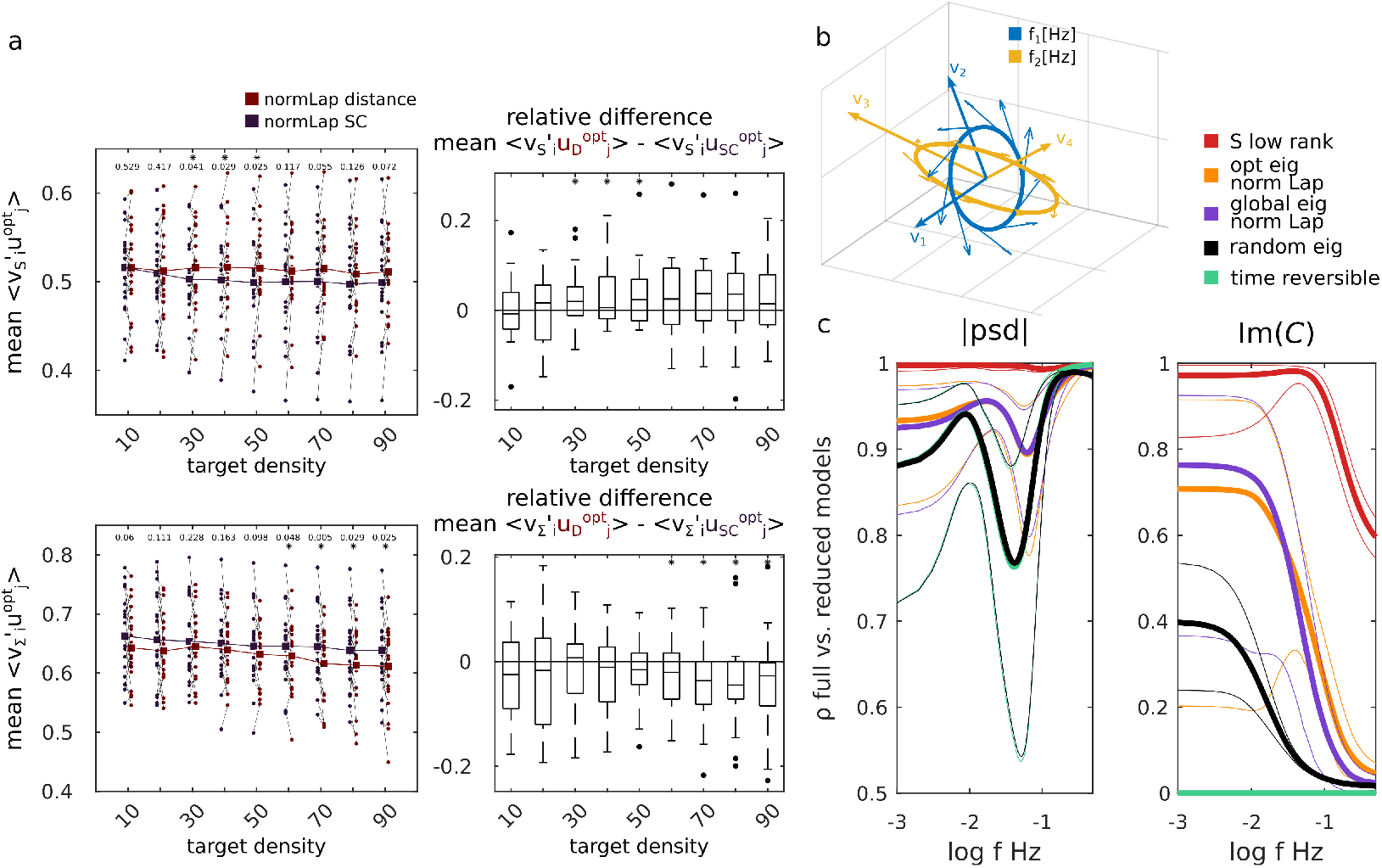
Alignment with structural connectivity and distance-based low-rank approximation. Panel *a*: for each mouse, the mean coupling between the dominant Schur modes and their optimally matched Laplacian eigenvectors, computed using the distance-based similarity matrix (red) and the structural connectivity matrix (blue) across different target densities (top left). The top right plot reports the relative difference between the two couplings, i.e. 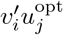, as a function of density. Bottom: the same analysis repeated using the dominant eigenvectors of each mouse-specific covariance matrix Σ. Panel *b*: schematic illustration of two orthogonal oscillatory modes. Panel *c*: absolute power spectral density |psd|, and imaginary part of coherency Im(*C*) reconstructed under different low-rank approximations of *S*: the two dominant Schur modes of *S* (red), the two dominant Schur pairs approximated using the corresponding optimally matched Laplacian eigenvectors (orange), the two dominant Schur pairs approximated using the four most frequently matched Laplacian eigenvectors across mice (blue), the two dominant Schur pairs constructed from four randomly generated orthogonal vectors (black), and the null model *S* = 0 (green).

We finally investigated whether the relationship between spatial distance and Schur modes could be exploited in a generative framework. Specifically, we asked whether the true *S* matrix could be approximated using distance-based Laplacian eigenvectors. To this end, we defined a gradient of approximation ranging from a low-rank representation of the true *S* matrix to the zero matrix, corresponding to time-reversible dynamics. Low-rank approximation of *S* were constructed by retaining the two dominant Schur mode pairs, preserving the subject-specific frequencies while varying the associated orthonormal bases (Figure 5b). We considered the following approximations (Figure 5c): (i) a low-rank reconstruction using the two dominant Schur modes (red); (ii) a reconstruction using the distance Laplacian eigenvectors optimally matched to the dominant Schur modes (orange); (iii) a reconstruction using the four distance Laplacian eigenvectors most frequently matched across mice (blue); (iv) a reconstruction using four randomly generated orthogonal vectors (black, average over 100 repetitions); and (v) the null model *S* = 0 (green). For each approximation, we evaluated the similarity between the true and reconstructed dynamics by comparing the absolute power spectral density matrices |psd|, and the imaginary part of coherency Im(*C*) over a grid of frequencies. We found that case (i) accurately reproduced both the absolute psd matrices and the Im(*C*), although a decrease in Im(*C*) correlation was observed at frequencies above 0.1 Hz. Cases (ii) and (iii), in which the dominant Schur modes were approximated using distance-based Laplacian eigenvectors, showed lower correlation than case (i); however, their performance was significantly better than the random approximation (case iv) within the frequency range of interest (0.01 - 0.1 Hz), corresponding to the empirical signal band. In contrast, case (v) largely overlapped with the random model (case iv) in reproducing the absolute psd, and exhibited zero correlation in Im(*C*), as expected. This in consistent with the fact that setting *S* = 0 yields time-reversible dynamics, for which the resulting psd is purely real.

## 4 Discussion

Top-down large-scale brain models are often criticized for their lack of interpretability. In contrast, bottom-up models adopt a more mechanistic generative approach that spans multiple scales in a forward manner. This work aims to reduce the gap between these two modeling paradigms. Within the top-down framework, linear state-space models constitute a standard formulation, where the Jacobian matrix encodes the so-called effective connectivity. Under this framework, when applied in the context of resting-state brain recordings, as the network size increases, we encounter not only practical challenges in model identification but also growing issues of interpretability.

The decomposition of the Jacobian matrix into the differential covariance and the precision matrix [35, 37], gives a more mechanistic view to the dynamical process [9]. In this work, we focused mainly on the role of differential covariance, and in particular on its antisymmetric component. Previous studies have shown that this antisymmetric term captures the rotational component of state-space trajectories and is directly related to entropy production, thus providing a quantitative measure of out-of-equilibrium dynamics [20, 25, 5].

At the macroscopic scale of brain activity, such rotational component, and the associated breaking of detailed balance, have received increasing attention in recent years [43]. These non-equilibrium signatures have been linked to task cognitive engagement, pathological conditions, and altered states of consciousness [32, 34, 38]. By leveraging the Schur decomposition, we analyzed the antisymmetric component of the differential covariance, obtaining a representation of the rotational dynamics as a linear combination of independent rotational modes. Each mode is characterized by a pair of orthogonal Schur basis vectors and an associated angular frequency. This angular frequency determines the rotational speed of circular trajectories in the corresponding state-space plane and, consequently, sets the phase lag between interacting signals. Based on this interpretation, we hypothesized that the spatial distance between brain regions plays a role in shaping the Schur basis on which these fixed-frequency rotation modes emerge.

This perspective resonates with what we refer to here as a bottom-up strategy, in which a specific local dynamical model is assigned to each network node and nodes are coupled according to the structural connectivity matrix [49]. In contrast to linear state-space models, where oscillatory components are expressed as global eigenmodes acting independently in the Schur space, bottom-up models assume node-specific oscillatory dynamics determined by the chosen parcellation. Recently, networks composed of oscillatory nodes have attracted increasing attention, not only as models of large-scale brain dynamics but also as a foundation for developing more efficient brain-inspired neural network [64, 18, 67, 13, 56].

Previous evidence indicates that spatial distance modulates the spectral properties of the brain signals and regulate inter-regional phase lags. Although these findings span different recording modalities and species, they consistently point to a frequency-dependent spatial organization. For instance, using ECoG data, in [42] authors showed that low-frequency activity remains correlated over large spatial distances than high-frequency activity. In the context of whole-brain modeling, an extension of the structural Laplacian incorporating distance-based time delay was proposed in [66]. This yielded a frequency-dependent complex Laplacian. The resulting eigenmodes showed improved alignment with canonical functional networks and revealed a clear frequency-specific organization. Complementary modeling studies have further demonstrated how distance-dependent delays shape large-scale synchrony patterns [69, 7]. In particular, these works showed that higher frequencies tend to promote local, intra-modular synchronization, whereas lower frequencies support long-range coordination. The preferential alignment we observed between faster Schur modes and the leading eigenvectors of the distance-based Laplacian suggests that spatial constraints selectively organize high-frequency rotational dynamics into geometrically local modes, while slower modes remain less constrained by distance. In this sense, the Schur decomposition of the antisymmetric part of the differential covariance provides a principled algebraic bridge between time-lagged dynamics, frequency-specific synchronization, and the spatial geometry of the brain.

However, the definition of the full Jacobian matrix critically depends on the precision matrix. Although our results highlight a relationship between the asymmetric component of the differential covariance and the spatial distance matrix, this should not be interpreted as a binding constraint, but rather as a first-order approximation of the oscillatory structure imposed by spatial embedding. The subsequent multiplication by the precision matrix mixes the Schur modes of the differential covariance and may break their independence by inducing a new Schur basis for the full Jacobian. A further limitation of relying solely on spatial distance is its inability to capture long-range connections. Although rare, such connections have been shown to play a fundamental role in shaping large-scale brain dynamics and are essential for reproducing empirical functional patterns [17].

The ability to approximate both the power spectral density and the imaginary part of coherency using only a few dominant Laplacian eigenvectors is consistent with the low dimensionality of the manifold on which large-scale brain dynamics evolves [16]. A substantial body of work has shown that resting-state brain activity can be effectively captured by low-dimensional graph-dynamical processes grounded into the structural Laplacian [2, 1, 63, 24]. These studies demonstrated that functional connectivity and structural Laplacian share similar eigenvectors. In this framework, resting-state networks can be reconstructed from the eigenvectors of the structural Laplacian, often referred to as connectome harmonics [3], and the degree of structure-functional coupling can be quantified at the node level using metrics such as the structural decoupling index [51].

Overall our results support a more mechanistic interpretation of linear state-space models. Within this framework, resting-state brain activity can be viewed as emerging from a set of independent rotational mode each defined by a characteristic frequency and phase lag. Interactions between these modes are mediated by the precision matrix, which governs their coupling and modulation.

Finally, this formulation opens new avenues for imposing anatomically grounded priors during model identification. By linking time-lagged dynamics to spatial distance constraints, it suggests principled ways to regularize the estimation of large-scale models, an issue that currently limits the applicability of effective connectivity approaches to fine parcellations. Incorporating geometrical information may therefore improve both interpretability and scalability of top-down models of resting-state brain dynamics.

## Acknowledgments

We thank S. Zampieri for fruitful discussions, as well as A. Gozzi and L. Coletta from the Functional Neuroimaging Laboratory (Center for Neuroscience and Cognitive Systems @ UniTn, Istituto Italiano di Tecnologia, Rovereto, Italy) for providing the mouse dataset and offering assistance in data curation and preprocessing.

## References

[1] F. Abdelnour, M. Dayan, O. Devinsky, T. Thesen, and A. Raj. Functional brain connectivity is predictable from anatomic network’s laplacian eigen-structure. NeuroImage, 172:728–739, 5 2018.

[2] F. Abdelnour, H. U. Voss, and A. Raj. Network diffusion accurately models the relationship between structural and functional brain connectivity networks. NeuroImage, 90, 2014.

[3] S. Atasoy, I. Donnelly, and J. Pearson. Human brain networks function in connectome-specific harmonic waves. Nature Communications 2016 7:1, 7:1–10, 1 2016.

[4] M. Belkin and P. Niyogi. Laplacian eigenmaps for dimensionality reduction and data representation. Neural Computation, 15:1373–1396, 6 2003.

[5] D. Benozzo, G. Baggio, G. Baron, A. Chiuso, S. Zampieri, and A. Bertoldo. Analyzing asymmetry in brain hierarchies with a linear state-space model of resting-state fmri data. Network Neuroscience, 8, 2024.

[6] D. Benozzo, G. Baron, L. Coletta, A. Chiuso, A. Gozzi, and A. Bertoldo. Macroscale coupling between structural and effective connectivity in the mouse brain. Scientific Reports, 14, 2024.

[7] E. Bolhasani, M. Hassanzadeh, M. Perc, and A. Valizadeh. Functional modularity induced by delayed interactions. Chaos, Solitons and Fractals, 198, 2025.

[8] T. Bolt, J. S. Nomi, D. Bzdok, J. A. Salas, C. Chang, B. T. T. Yeo, L. Q. Uddin, and S. D. Keilholz. A parsimonious description of global functional brain organization in three spatiotemporal patterns. Nature Neuroscience, 25:1093–1103, 8 2022.

[9] U. Casti, G. Baggio, D. Benozzo, S. Zampieri, A. Bertoldo, and A. Chiuso. Dynamic brain networks with prescribed functional connectivity. Proceedings of the IEEE Conference on Decision and Control, pages 709–714, 2023.

[10] Y. Chen, Q. Bukhari, T. W. Lin, and T. J. Sejnowski. Functional connectivity of fmri using differential covariance predicts structural connectivity and behavioral reaction times. Network Neuroscience, 6:614–633, 6 2022.

[11] T. B. Clark and A. D. Maestro. Moments of the inverse participation ratio for the laplacian on finite regular graphs. Journal of Physics A: Mathematical and Theoretical, 51:495003, 11 2018.

[12] L. D. Costa, K. Friston, C. Heins, and G. A. Pavliotis. Bayesian mechanics for stationary processes. Proceedings of the Royal Society A: Mathematical, Physical and Engineering Sciences, 477, 12 2021.

[13] T. Dan, J. Ding, and G. Wu. Explore brain-inspired machine intelligence for connecting dots on graphs through holographic blueprint of oscillatory synchronization. Nature Communications, 16, 2025.

[14] E. D’Angelo and V. Jirsa. The quest for multiscale brain modeling. Trends in neurosciences, 45:777–790, 10 2022.

[15] G. Deco and M. L. Kringelbach. Turbulent-like dynamics in the human brain. Cell Reports, 33, 2020.

[16] G. Deco, Y. S. Perl, and M. L. Kringelbach. Complex harmonics reveal low-dimensional manifolds of critical brain dynamics. Physical Review E, 111, 2025.

[17] G. Deco, Y. S. Perl, P. Vuust, E. Tagliazucchi, H. Kennedy, and M. L. Kringelbach. Rare long-range cortical connections enhance human information processing. Current Biology, 31, 2021.

[18] F. Effenberger, P. Carvalho, I. Dubinin, and W. Singer. The functional role of oscillatory dynamics in neocortical circuits: A computational perspective. Proceedings of the National Academy of Sciences of the United States of America, 122, 2025.

[19] M. Ercsey-Ravasz, N. T. Markov, C. Lamy, D. C. VanEssen, K. Knoblauch, Z. Toroczkai, and H. Kennedy. A predictive network model of cerebral cortical connectivity based on a distance rule. Neuron, 80, 2013.

[20] K. J. Friston, E. D. Fagerholm, T. S. Zarghami, T. Parr, I. Hipólito, L. Magrou, and A. Razi. Parcels and particles: Markov blankets in the brain. Network Neuroscience, 5:211–251, 2 2021.

[21] K. J. Friston, L. Harrison, and W. Penny. Dynamic causal modelling. NeuroImage, 19:1273–1302, 8 2003.

[22] K. J. Friston, A. Mechelli, R. Turner, and C. J. Price. Nonlinear responses in fmri: The balloon model, volterra kernels, and other hemodynamics. NeuroImage, 12, 2000.

[23] S. Frässle, E. I. Lomakina, A. Razi, K. J. Friston, J. M. Buhmann, and K. E. Stephan. Regression dcm for fmri. NeuroImage, 3 2017.

[24] S. Ghosh, A. Raj, and S. S. Nagarajan. A joint subspace mapping between structural and functional brain connectomes. NeuroImage, 272, 2023.

[25] M. Gilson, E. Tagliazucchi, and R. Cofré. Entropy production of multivariate ornstein-uhlenbeck processes correlates with consciousness levels in the human brain. Physical Review E, 107:024121, 2 2023.

[26] Y. Gu, L. E. Sainburg, S. Kuang, F. Han, J. W. Williams, Y. Liu, N. Zhang, X. Zhang, D. A. Leopold, and X. Liu. Brain activity fluctuations propagate as waves traversing the cortical hierarchy. Cerebral Cortex, 31, 2021.

[27] D. Gutierrez-Barragan, M. A. Basson, S. Panzeri, and A. Gozzi. Infraslow state fluctuations govern spontaneous fmri network dynamics. Current Biology, 29, 2019.

[28] D. Gutierrez-Barragan, N. A. Singh, F. G. Alvino, L. Coletta, F. Rocchi, E. D. Guzman, A. Galbusera, M. Uboldi, S. Panzeri, and A. Gozzi. Unique spatiotemporal fmri dynamics in the awake mouse brain. Current biology : CB, 32:631–644.e6, 2 2022.

[29] R. Gămănuţ, H. Kennedy, Z. Toroczkai, M. Ercsey-Ravasz, D. C. V. Essen, K. Knoblauch, and A. Burkhalter. The mouse cortical connectome, characterized by an ultra-dense cortical graph, maintains specificity by distinct connectivity profiles. Neuron, 97, 2018.

[30] S. Horvát, R. Gămănut,, M. Ercsey-Ravasz, L. Magrou, B. Gămănut,, D. C. V. Essen, A. Burkhalter, K. Knoblauch, Z. Toroczkai, and H. Kennedy. Spatial embedding and wiring cost constrain the functional layout of the cortical network of rodents and primates. PLoS Biology, 14, 2016.

[31] Y. Hosaka, T. Hieda, K. Hayashi, K. Jimura, and T. Matsui. Linear models replicate the energy landscape and dynamics of resting-state brain activity. bioRxiv, 2024.

[32] S. Idesis, M. Allegra, J. Vohryzek, Y. S. Perl, J. Faskowitz, O. Sporns, M. Corbetta, and G. Deco. A low dimensional embedding of brain dynamics enhances diagnostic accuracy and behavioral prediction in stroke. Scientific Reports 2023 13:1, 13:1–17, 9 2023.

[33] M. Józsa, M. Ercsey-Ravasz, and Z. I. Lázár. Coarse-graining model reveals universal exponential scaling in axonal length distributions. Journal of Physics: Complexity, 5, 2024.

[34] M. L. Kringelbach, Y. S. Perl, E. Tagliazucchi, and G. Deco. Toward naturalistic neuroscience: Mechanisms underlying the flattening of brain hierarchy in movie-watching compared to rest and task. Science Advances, 9, 1 2023.

[35] C. Kwon, P. Ao, and D. J. Thouless. Structure of stochastic dynamics near fixed points. Proceedings of the National Academy of Sciences, 102:13029–13033, 9 2005.

[36] R. Lambiotte and M. T. Schaub. Modularity and dynamics on complex networks. Modularity and Dynamics on Complex Networks, 2 2021.

[37] T. W. Lin, A. Das, G. P. Krishnan, M. Bazhenov, and T. J. Sejnowski. Differential covariance: A new class of methods to estimate sparse connectivity from neural recordings. Neural Computation, 29:2581–2632, 10 2017.

[38] C. W. Lynn, E. J. Cornblath, L. Papadopoulos, M. A. Bertolero, and D. S. Bassett. Broken detailed balance and entropy production in the human brain. Proceedings of the National Academy of Sciences of the United States of America, 118:e2109889118, 11 2021.

[39] M. Mancini, Q. Tian, Q. Fan, M. Cercignani, and S. Y. Huang. Dissecting whole-brain conduction delays through mri microstructural measures. Brain Structure and Function, 226, 2021.

[40] T. Matsui, T. Q. Pham, K. Jimura, and J. Chikazoe. On co-activation pattern analysis and non-stationarity of resting brain activity. NeuroImage, 249:118904, 4 2022.

[41] A. Mitra, A. Z. Snyder, C. D. Hacker, and M. E. Raichle. Lag structure in resting-state fmri. Journal of Neurophysiology, 111, 2014.

[42] L. Muller, L. S. Hamilton, E. Edwards, K. E. Bouchard, and E. F. Chang. Spatial resolution dependence on spectral frequency in human speech cortex electrocorticography. Journal of Neural Engineering, 13, 2016.

[43] R. Nartallo-Kaluarachchi, M. Kringelbach, G. Deco, R. Lambiotte, and A. Goriely. Nonequilibrium physics of brain dynamics, 2026.

[44] G. Nolte, O. Bai, L. Wheaton, Z. Mari, S. Vorbach, and M. Hallett. Identifying true brain interaction from eeg data using the imaginary part of coherency. Clinical neurophysiology, 115:2292–2307, 10 2004.

[45] E. Nozari, M. A. Bertolero, J. Stiso, L. Caciagli, E. J. Cornblath, X. He, A. S. Mahadevan, G. J. Pappas, and D. S. Bassett. Macroscopic resting-state brain dynamics are best described by linear models. Nature Biomedical Engineering 2023 8:1, 8:68–84, 12 2023.

[46] S. W. Oh, J. A. Harris, L. Ng, B. Winslow, N. Cain, S. Mihalas, Q. Wang, C. Lau, L. Kuan, A. M. Henry, M. T. Mortrud, B. Ouellette, T. N. Nguyen, S. A. Sorensen, C. R. Slaughterbeck, W. Wakeman, Y. Li, D. Feng, A. Ho, E. Nicholas, K. E. Hirokawa, P. Bohn, K. M. Joines, H. Peng, M. J. Hawrylycz, J. W. Phillips, J. G. Hohmann, P. Wohnoutka, C. R. Gerfen, C. Koch, A. Bernard, C. Dang, A. R. Jones, and H. Zeng. A mesoscale connectome of the mouse brain. Nature 2014 508:7495, 508:207–214, 4 2014.

[47] J. C. Pang, K. M. Aquino, M. Oldehinkel, P. A. Robinson, B. D. Fulcher, M. Breakspear, and A. Fornito. Geometric constraints on human brain function. Nature, 618, 2023.

[48] Y. S. Perl, H. Bocaccio, C. Pallavicini, I. Pérez-Ipiña, S. Laureys, H. Laufs, M. Kringelbach, G. Deco, and E. Tagliazucchi. Nonequilibrium brain dynamics as a signature of consciousness. Physical Review E, 104:014411, 7 2021.

[49] A. Ponce-Alvarez and G. Deco. The hopf whole-brain model and its linear approximation. Scientific Reports 2024 14:1, 14:2615–, 1 2024.

[50] G. Prando, M. Zorzi, A. Bertoldo, M. Corbetta, M. Zorzi, and A. Chiuso. Sparse dcm for whole-brain effective connectivity from resting-state fmri data. NeuroImage, 208, 2020.

[51] M. G. Preti and D. V. D. Ville. Decoupling of brain function from structure reveals regional behavioral specialization in humans. Nature Communications 2019 10:1, 10:1–7, 10 2019.

[52] B. Péntek and M. Ercsey-Ravasz. The exponential distance rule-based network model predicts topology and reveals functionally relevant properties of the drosophila projectome. Network Neuroscience, 9, 2025.

[53] R. V. Raut, A. Z. Snyder, A. Mitra, D. Yellin, N. Fujii, R. Malach, and M. E. Raichle. Global waves synchronize the brain’s functional systems with fluctuating arousal. Science Advances, 7, 2021.

[54] A. Razi, M. L. Seghier, Y. Zhou, P. McColgan, P. Zeidman, H. J. Park, O. Sporns, G. Rees, and K. J. Friston. Large-scale dcms for resting-state fmri. Network neuroscience (Cambridge, Mass.), 1:222–241, 1 2017.

[55] F. Rocchi, C. Canella, S. Noei, D. Gutierrez-Barragan, L. Coletta, A. Galbusera, A. Stuefer, S. Vassanelli, M. Pasqualetti, G. Iurilli, S. Panzeri, and A. Gozzi. Increased fmri connectivity upon chemogenetic inhibition of the mouse prefrontal cortex. Nature Communications 2022 13:1, 13:1–15, 2 2022.

[56] N. R. Rohan, C. Vigneswaran, S. Ghosh, K. Rajendran, A. Gaurav, and V. S. Chakravarthy. Deep oscillatory neural network. Scientific Reports, 15, 2025.

[57] F. E. Rosas, A. I. Luppi, P. A. Mediano, M. L. Kringelbach, L. Pessoa, and F. Turkheimer. Top-down and bottom-up neuroscience: overcoming the clash of research cultures, 2025.

[58] R. Salvador, J. Suckling, M. R. Coleman, J. D. Pickard, D. Menon, and E. Bullmore. Neurophysiological architecture of functional magnetic resonance images of human brain. Cerebral Cortex, 15, 2005.

[59] R. Salvador, J. Suckling, C. Schwarzbauer, and E. Bullmore. Undirected graphs of frequency-dependent functional connectivity in whole brain networks. Philosophical Transactions of the Royal Society B: Biological Sciences, 360, 2005.

[60] J. Shi and J. Malik. Normalized cuts and image segmentation. IEEE Transactions on Pattern Analysis and Machine Intelligence, 22:888–905, 2000.

[61] M. Shinn, A. Hu, L. Turner, S. Noble, K. H. Preller, J. L. Ji, F. Moujaes, S. Achard, D. Scheinost, R. T. Constable, J. H. Krystal, F. X. Vollenweider, D. Lee, A. Anticevic, E. T. Bullmore, and J. D. Murray. Functional brain networks reflect spatial and temporal autocorrelation. Nature Neuroscience 2023 26:5, 26:867–878, 4 2023.

[62] B. S. Sipes, S. S. Nagarajan, and A. Raj. Integrative, segregative, and degenerate harmonics of the structural connectome. Communications Biology 2024 7:1, 7:986–, 8 2024.

[63] P. Tewarie, B. Prasse, J. M. Meier, F. A. Santos, L. Douw, M. M. Schoonheim, C. J. Stam, P. V. Mieghem, and A. Hillebrand. Mapping functional brain networks from the structural connectome: Relating the series expansion and eigenmode approaches. NeuroImage, 216, 2020.

[64] A. Todri-Sanial, C. Delacour, M. Abernot, and F. Sabo. Computing with oscillators from theoretical underpinnings to applications and demonstrators. npj Unconventional Computing, 1, 2024.

[65] Q. Wang, S.-L. Ding, Y. Li, J. Royall, D. Feng, P. Lesnar, N. Graddis, M. Naeemi, B. Facer, A. Ho, T. Dolbeare, B. Blanchard, N. Dee, W. Wakeman, K. E. Hirokawa, A. Szafer, S. M. Sunkin, S. W. Oh, A. Bernard, J. W. Phillips, M. Hawrylycz, C. Koch, H. Zeng, J. A. Harris, and L. Ng. The allen mouse brain common coordinate framework: A 3d reference atlas. Cell, 181:936–953.e20, 5 2020.

[66] X. Xie, C. Cai, P. F. Damasceno, S. S. Nagarajan, and A. Raj. Emergence of canonical functional networks from the structural connectome. NeuroImage, 237, 2021.

[67] Y. Yan, Q. Yang, Y. Wu, H. Liu, M. Zhang, H. Li, K. C. Tan, and J. Wu. Efficient and robust temporal processing with neural oscillations modulated spiking neural networks. Nature Communications, 16, 2025.

[68] B. Yousefi and S. Keilholz. Propagating patterns of intrinsic activity along macroscale gradients coordinate functional connections across the whole brain. NeuroImage, 231, 2021.

[69] A. Ziaeemehr, M. Zarei, A. Valizadeh, and C. R. Mirasso. Frequency-dependent organization of the brain’s functional network through delayed-interactions. Neural Networks, 132, 2020.

